# Genome-wide characterization of the R2R3-MYB transcription factors in pepper (*Capsicum* spp.) unveils the role of CaMYB101 as repressor in anthocyanin biosynthesis

**DOI:** 10.1101/2021.08.24.457473

**Authors:** Ying Liu, Yi Wu, Zicheng Wang, Shiya Zhang, Xintong Liu, Yury Tikunov, Richard G.F. Visser, Rob E. Schouten, Leo F.M. Marcelis, Zhao Zhang, Arnaud Bovy

**Affiliations:** Plant Breeding, Wageningen University and Research, Wageningen, Netherlands; Horticulture and Product Physiology, Wageningen University and Research, Wageningen, Netherlands; Graduate School Production Ecology & Resource Conservation, Wageningen University and Research, Wageningen, Netherlands; Graduate School Experimental Plant Sciences, Wageningen University and Research, Wageningen, the Netherlands; Beijing Key Laboratory of Development and Quality Control of Ornamental Crops, Department of Ornamental Horticulture, China Agricultural University, Beijing, China

**Keywords:** pepper, MYB transcription factor, *Capsicum annuum*, *Capsicum chinense*, *Capsicum baccatum*

## Abstract

Fruit colour is one of the most important commercial traits of pepper (*Capsicum* spp.), a major horticultural crop worldwide. Some pepper accessions temporarily accumulate anthocyanins during fruit development and gradually lose them upon fruit ripening. Meanwhile, anthocyanin biosynthesis gradually stops. However, how this process is exactly regulated is still largely unknown. R2R3-MYB is one of the largest plant transcription factor families, and it is considered the most important regulator for the biosynthesis of anthocyanins and other flavonoids. Although R2R3-MYBs are widely studied in many plants, research in pepper has been limited. In this study, we performed a genome-wide analysis of R2R3-MYBs across three cultivated pepper species (*C. annuum*, *C. baccatum*, and *C. chinense* ) involving identification, chromosome localization, gene structure analysis, phylogenetic analysis and collinearity analysis. Candidate R2R3-MYB repressors were further identified based on repression motifs. An R2R3-MYB gene, *CaMYB101*, was selected based on its high homology with anthocyanin biosynthesis repressors in tomato and petunia as well as its high expression level in fruit when purple pigmentation started to discolour. By using virus-induced gene silencing, CaMYB101 was characterized as an anthocyanin biosynthesis repressor. To our knowledge, CaMYB101 is the first transcriptional repressor associated with anthocyanin biosynthesis identified in pepper.

## Introduction

Transcription factors (TFs) are involved in stimulating or repressing the transcription of target genes to control physiological and metabolic processes during plant growth and development in response to endogenous or exogenous stimuli (Fuda et al., 2009). Based on their DNA-binding domains, TFs can be classified into different families. The MYB family is one of the largest TF families in plants and is involved in various biological processes including plant growth, circadian clock control, and primary and secondary metabolism regulation (Dubos et al., 2010). A highly conserved DNA-binding domain, known as the MYB domain, contains up to four incomplete repeats at the N-terminus. Each repeat is composed of approximately 50-55 amino acid residues and forms three α-helices. The second and third helices form a helix-turn-helix architecture with three spaced tryptophan (or hydrophobic) residues (Ogata et al., 1992; Du et al., 2015). The C-terminus region of MYB proteins is highly variable and responsible for the diverse regulatory functions of MYB TFs. Depending on the number of tandem repeat(s) in the MYB domain, the MYB family has four subfamilies: 1R-MYB (MYB-related), 2R-MYB (R2R3-MYB), 3R-MYB (R1R2R3-MYB) and 4R-MYB (four R1/R2-like repeats) (Dubos et al., 2010). In plants, the first MYB (R2R3-MYB) TF COLORED1 (C1) was discovered from *Zea mays* and has been identified to be involved in the activation of anthocyanin biosynthesis (Grotewold et al. 1991). Over time, the MYB family has been identified in many plant species. In *Arabidopsis*, 198 AtMYBs were identified and the tomato genome included 127 SlMYBs (Yanhui et al., 2006; Li et al., 2016). Among the MYB subfamilies, R2R3-MYB is the dominant subfamily with various roles in, for example, flavonoid biosynthesis.

Flavonoids are a large group of secondary metabolites, including flavones, isoflavones, flavonols, anthocyanins and proanthocyanidins. The various flavonoids can serve as antioxidants, defense compounds, signaling molecules and pigments. In plants, there are two types of structural genes: (i) early flavonoid biosynthesis pathway genes (EBGs), encoding chalcone synthase (CHS), chalcone isomerase (CHI), flavonol 3-hydroxylase (F3H) and flavonol 3’-hydroxylase (F3’H), which are induced as precursors for the downstream genes, known as late biosynthetic genes (LBGs). (ii) The LBGs including dihydroflavonol 4-reductase (DFR), anthocyanin synthase (ANS) and UDP-glucose: flavonoid 3-glucosyltransferase (UFGT), leucoanthocyanidin reductase (LAR) and anthocyanin reductase (ANR) which lead to proanthocyanidin and anthocyanin biosynthesis. The regulation of flavonoid biosynthetic structural genes often involves R2R3-MYB TFs. The EGBs are activated by independent R2R3-MYBs such as AtMYB11, AtMYB12 and AtMYB111, while LBGs related with anthocyanin biosynthesis are regulated by a MYB-bHLH-WD40 (MBW) complex consisting of a MYB TF, a basic helix-loop-helix (bHLH) and a WD-repeat protein (Xu et al., 2015). There is one exception, F3’H, which is regulated by both independent R2R3-MYBs and an MBW complex (Stracke et al., 2007).

To date, the majority of the flavonoid biosynthesis R2R3-MYBs are transcriptional activators. For example, high anthocyanin pigmented apple fruits were observed by overexpression of apple *MdMYB10* in cv. ‘Royal Gala’, together with increased expression of *MdCHS*, *MdCHI*, *MdF3H*, *MdDFR*, *MdANS* and *MdUFGT* (Espley et al., 2007). In grapevine, VvMYB5a leads to an up-regulation of flavonoid biosynthesis and boosted flavonols, anthocyanins and proanthocyanins accumulation (Deluc et al., 2006; Deluc et al., 2008). In addition to flavonoid activators, R2R3-MYB repressors have also been identified and displayed a crucial role in balancing flavonoid biosynthesis. For instance, the grapevine VvMYB4-like and strawberry FaMYB1 suppress the flavonol and anthocyanin accumulation and cause a clear pigmentation loss in flowers (Aharoni et al., 2001; Perez-Diaz et al., 2016). In petunia, overexpression of *PhMYB27* causes reduced anthocyanin accumulation, while silencing of this gene enhances anthocyanin accumulation throughout the whole plant including leaves, stems, and flowers (Albert et al., 2014).

Pepper is one of the most popular horticultural crops worldwide. The cultivated pepper comprises several species within the genus *Capsicum*, such as *C. annuum*, *C. baccatum* and *C. chinense*. The fruits of pepper contain a wide range of health-related secondary metabolites, like carotenoids and flavonoids. Among peppers, there are some purple accessions with a high level of anthocyanins in fruit peels. Moreover, the overall flavonoid content of these purple accessions is generally higher compared to non-purple accessions (Liu et al., 2020). These purple pigments are due to a temporary anthocyanin accumulation before fruit ripening and after ripening the anthocyanin biosynthesis stops and degradation becomes dominant (Yamada et al., 2019). Several R2R3-MYBs have been reported to activate flavonoid biosynthesis in pepper (Borovsky et al., 2004; Zhang et al., 2015). However, no MYB repressors for flavonoid biosynthesis have been identified in pepper yet. Genome-wide identification of the R2R3-MYB gene family has been studied in *C. annuum* (Wang et al., 2020; Arce-Rodríguez et al., 2021). The recently released pepper genomes make the study of the R2R3-MYB gene family in *C. baccatum* and *C. chinense* possible. We aim to globally analyse the R2R3-MYB gene family in three *Capsicum* species and among them identify flavonoid-related R2R3-MYB represosrs in pepper. To accomplish this, a genome-wide analysis of R2R3-MYBs was performed within the three cultivated *Capsicum* species (*C. annuum*, *C. baccatum* and *C. chinense*), including gene structure, chromosomal location, synteny and phylogeny. As the result, candidate R2R3-MYB repressors in *Capsicum* spp. were identified. We showed that a R2R3-MYB repressor, named CaMYB101, plays a role in the negative regulation of anthocyanin biosynthesis in pepper. Our results lay a foundation for characterizing R2R3-MYBs among *Capsicum* spp. and provide a better understanding for flavonoid-related repression mechanism in pepper.

## Results

### Identification of R2R3-MYBs in *Capsicum* spp. genomes

The published *C. baccatum* and *C. chinense* genome sequences (Table 1) were used for putative R2R3-MYBs identification through HMM profiling (PF00249). After removing redundant proteins by BLASTP through NCBI-CDD, 106 and 110 R2R3-MYB transcription factors were identified in *C. baccatum* (CbMYBs) and *C. chinense* (CcMYBs), respectively. Meanwhile, 108 R2R3-MYBs of *C. annuum* (CaMYBs) were derived from the study of Wang et al. (2020). All R2R3-MYBs were named based on their species and chromosomal orders, with details including the accession number, chromosomal location, number of introns and exons as well as the length of coding sequence and protein (Table S1-S3). The longest CaMYB is CaMYB82 with 995 amino acids and the shortest is CaMYB58 with 110 amino acids. CbMYB21 is the longest MYB protein of all *Capsicum* spp., and contains 1210 amino acids, while CbMYB10 is the shortest CbMYB protein with 152 amino acids. The length of CcMYB proteins varied from 137 (CcMYB66) to 1156 (CcMYB42) amino acids.

**Table 1.** Assembly descriptions and sources for three genomes used to characterize R2R3-MYB diversity in *Capsicum* spp.

| Genome features | *Capsicum annuum*<sup>1</sup> | *Capsicum baccatum*<sup>2</sup> | *Capsicum chinense*<sup>2</sup> |
| --- | --- | --- | --- |
| Assembled genome size (Gb) | 2.9 | 3.2 | 3.0 |
| Number of Scaffolds | 6478 | 2083 | 1557 |
| Scaffold N50 (Mb) | 1.4 | 2.0 Mb | 3.3 Mb |
| Coverage | 99X | 98X | 80X |

<sup>1</sup> Pepper Zunla 1 Ref\_v1.0 (https://www.ncbi.nlm.nih.gov/assembly/GCA_000710875.1);
<sup>2</sup> Kim et al. (2017).

### Sequence conservation within the R2 and R3 domains

Multiple sequence alignment analyses of all putative R2R3-MYBs derived from three *Capsicum* spp. as well as from *Arabidopsis* were performed to demonstrate the feature of homologous domain sequences and the corresponding amino acid frequency. Similar to *Arabidopsis*, the R2 and R3 domains of all the R2R3-MYBs among *Capsicum* spp. were highly conserved (Figure 1). Meanwhile, the tryptophans (W) are highly conserved in R2 (three) and R3(two) domains.

**Figure 1.**
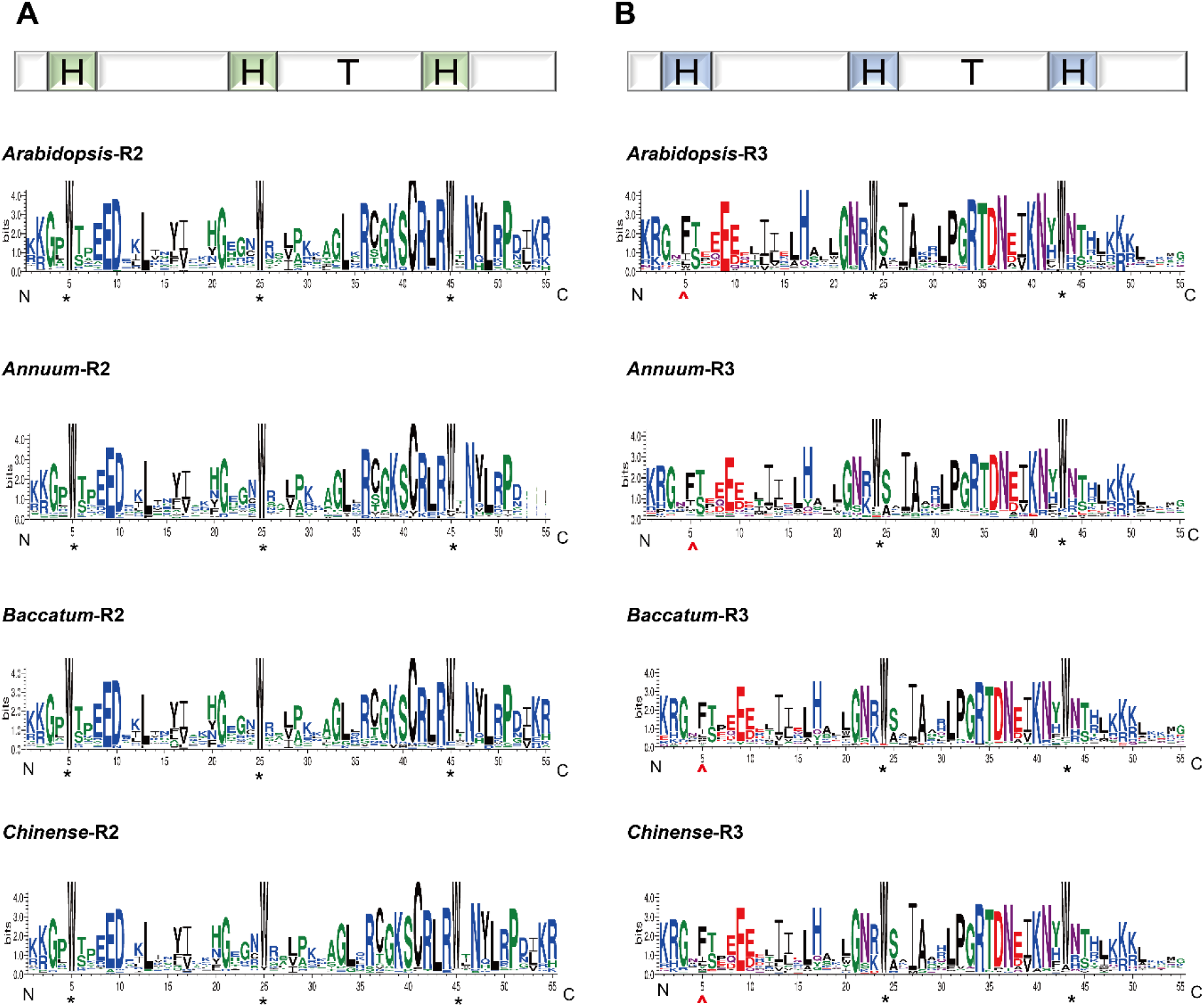
The conserved motifs of the R2 and R3 domains in *Capsicum* spp. R2R3-MYB proteins. (A) The R2 domain of *C. annuum, C. baccatum and C. chinense;* (B) the R3 domain of *C. annuum, C. baccatum* and *C. chinense*. These sequence logos were determined from the multiple alignment analysis of 108, 106 and 110 R2R3-MYB proteins in *C. annuum, C. baccatum* and *C. chinense* respectively. Each MYB repeat contains three α-helices (H). The second and third helices form a helix-turn-helix architecture (HTH). The bit score shows the information content for each position in the sequence. The conserved tryptophan residues (W) are marked with black asterisks and the replacement of tryptophan in the R3 repeat are marked by red circumflex accent.

### Chromosomal localization and collinearity/ synteny analysis of *Capsicum* R2R3-MYBs

The Ca-, Cb- and CcMYBs were physically mapped throughout all 12 chromosomes in *C. annuum*, *C. baccatum* and *C. chinense*, respectively (Figure 2). In total 11 CaR2R3-MYBs, 9 CbR2R3-MYBs and 5 CcR2R3-MYBs were located on unanchored scaffolds and assigned to Chr. 00 (Table S1-S3). The distribution of R2R3-MYBs among three *Capsicum* spp. genomes was also not congruent. *C. annuum* and *C. chinense* had a relatively high density of R2R3-MYBs at the top and bottom of chromosomes compared to *C. baccatum*. The maximum and minimum number of R2R3-MYBs allocated per genome were similar. In *C. annuum*, Chr. 03 contained the most CaR2R3-MYBs, 12, whereas Chr. 08 had the lowest number of CaR2R3-MYBs, four, over all chromosomes (Figure 2A). In *C. baccatum*, Chr. 01, 03, 05, 06 and 07 contained the highest number (10) CbR2R3-MYBs, while Chr. 04 and 09 had the lowest number of four CbR2R3-MYBs (Figure 2B). In *C. chinense*, Chr. 06 had the highest number (13) CcR2R3-MYBs while Chr. 08 contained only two CcR2R3-MYBs (Figure 2C).

**Figure 2.**
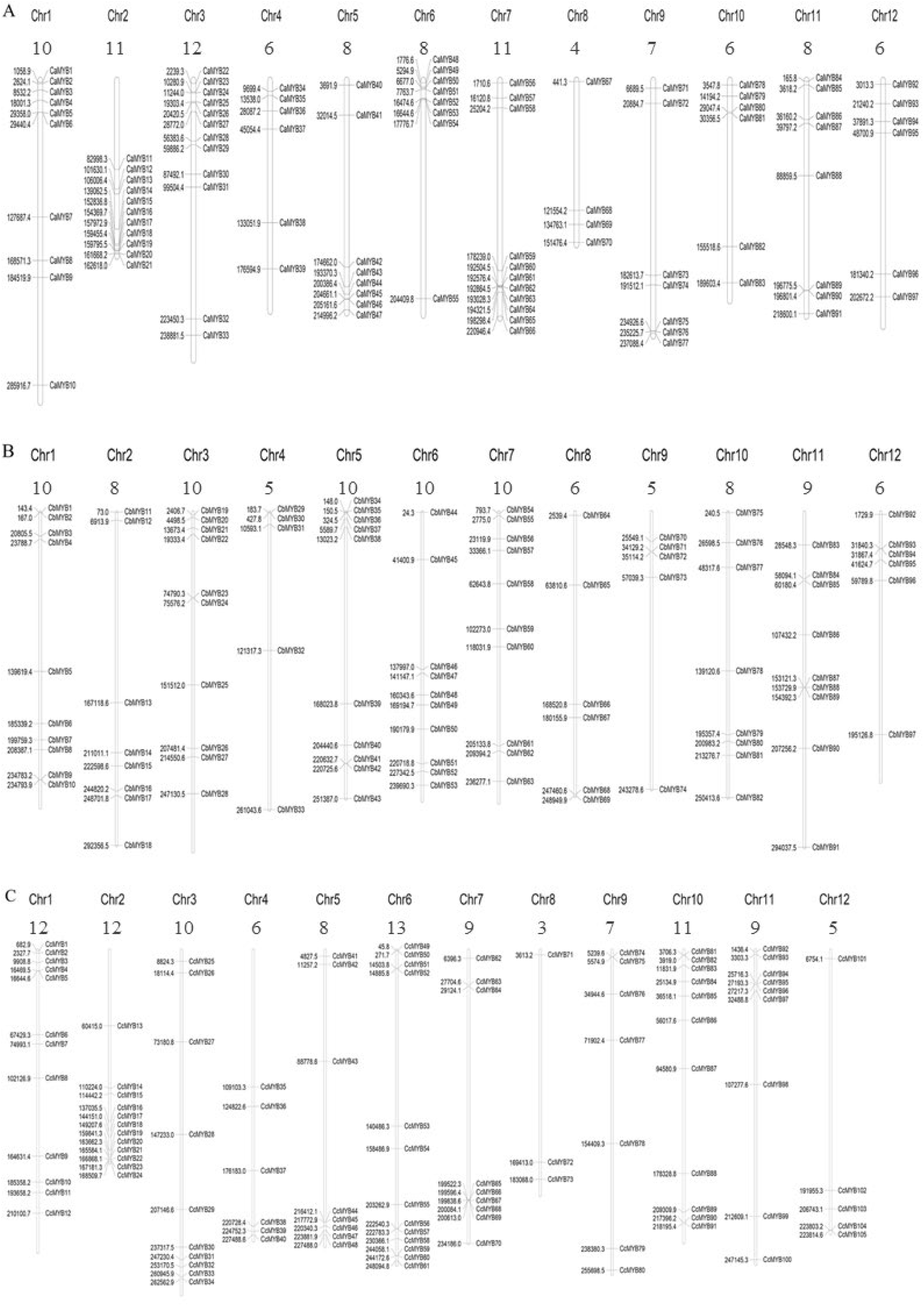
Chromosome localization of *Capsicum* spp. R2R3-MYB family members. The physical distribution of each R2R3-MYB gene is listed on the chromosomes of (A) *Capsicum annuum*, (B) *Capsicum baccatum* and (C) *Capsicum chinense*.

In total, 119 syntenic gene pairs were identified (Table 2, Supplemental Table S4), including 95 pairs of inter-species orthologs and 24 pairs of intra-species paralogs. To be specific, 44 gene pairs were between *C. annuum* and *C. chinense*, 30 gene pairs were between *C. annuum* and *C. baccatum* and 21 gene pairs were between *C. baccatum* and *C. chinense* (Figure 3, Supplemental Table S4). Among the 24 syntenic paralogs, 21 gene pairs were inter-chromosomal, while the rest three pairs were located on chromosome 2 (CaMYB12-CaMYB19, CaMYB13-CaMYB20 and CcMYB14-CcMYB22). The synonymous (Ks) and non-synonymous (Ka) values were used to determine the selection pressures. The Ka/Ks ratios for segmental duplication were between 0.11 and 0.53 through syntenic paralogs. In the syntenic orthologs, two out of 95 pairs showed Ka/Ks ratios larger than 1.0, indicating a positive selection on these gene pairs (for improved function), while the rest 93 pairs harboured the Ka/Ks smaller than 1.0, indicating stabilising selection (to maintain function).

**Table 2.** Ka-Ks calculation of each pair of syntenic R2R3-MYB paralogs within *Capsicum annuum*, *Capsicum baccatum* and *Capsicum chinense*, respectively. The minimum block size for collinear analysis is set at 5 in MCSanX for intraspecies analysis.

| Sequence1 | Sequence 2 | Ka<sup>1</sup> | Ks<sup>2</sup> | Ka/Ks | Effective Len<sup>3</sup> | Average S-sites<sup>4</sup> | Average N-sites<sup>5</sup> |
| --- | --- | --- | --- | --- | --- | --- | --- |
| CaMYB12 | CaMYB19 | 0.20 | 0.84 | 0.24 | 858 | 186.67 | 671.33 |
| CaMYB13 | CaMYB20 | 0.13 | 0.63 | 0.20 | 585 | 120.67 | 464.33 |
| CaMYB19 | CaMYB24 | 0.34 | 2.54 | 0.13 | 864 | 190.33 | 673.67 |
| CaMYB16 | CaMYB34 | 0.27 | 1.44 | 0.18 | 783 | 190.42 | 592.58 |
| CaMYB15 | CaMYB35 | 0.46 | 1.56 | 0.29 | 864 | 186.42 | 677.58 |
| CaMYB12 | CaMYB54 | 0.33 | 1.47 | 0.23 | 957 | 209.58 | 747.42 |
| CaMYB16 | CaMYB93 | 0.34 | 1.72 | 0.20 | 636 | 145.67 | 490.33 |
| CaMYB26 | CaMYB52 | 0.22 | 0.43 | 0.50 | 867 | 176.58 | 690.42 |
| CaMYB23 | CaMYB91 | 0.37 | 3.55 | 0.11 | 765 | 154.25 | 610.75 |
| CaMYB34 | CaMYB93 | 0.26 | 1.71 | 0.15 | 726 | 162.67 | 563.33 |
| CaMYB51 | CaMYB85 | 0.31 | 0.82 | 0.38 | 858 | 173.83 | 684.17 |
| CaMYB64 | CaMYB79 | 0.15 | 0.80 | 0.19 | 942 | 199.08 | 742.92 |
| CaMYB76 | CaMYB83 | 0.14 | 0.75 | 0.19 | 381 | 79.25 | 301.75 |
| CbMYB106 | CbMYB13 | 0.25 | 1.41 | 0.18 | 867 | 206.17 | 660.83 |
| CbMYB12 | CbMYB29 | 0.44 | 1.91 | 0.23 | 858 | 186.08 | 671.92 |
| CcMYB107 | CcMYB14 | 0.33 | 1.53 | 0.22 | 957 | 208.58 | 748.42 |
| CcMYB12 | CcMYB50 | 0.33 | 0.83 | 0.39 | 807 | 162.25 | 644.75 |
| CcMYB14 | CcMYB22 | 0.20 | 0.89 | 0.23 | 858 | 186.67 | 671.33 |
| CcMYB16 | CcMYB39 | 0.25 | 1.42 | 0.17 | 855 | 203.17 | 651.83 |
| CcMYB19 | CcMYB38 | 0.40 | 1.70 | 0.24 | 855 | 185.08 | 669.92 |
| CcMYB31 | CcMYB56 | 0.23 | 0.44 | 0.53 | 882 | 179.67 | 702.33 |
| CcMYB40 | CcMYB103 | 0.51 | 2.17 | 0.24 | 615 | 140.25 | 474.75 |
| CcMYB39 | CcMYB103 | 0.26 | 1.70 | 0.15 | 726 | 163.83 | 562.17 |
| CcMYB41 | CcMYB101 | 0.26 | 0.91 | 0.29 | 849 | 170.25 | 678.75 |

<sup>1</sup> Ka, non-synonymous substitution rate;
<sup>2</sup> Ks, synonymous substitution rate;
<sup>3</sup> Effective Len: effective length of compared sequences;
<sup>4</sup> S-site, number of synonymous site;
<sup>5</sup> N-site, number of non-synonymous sites.

**Figure 3.**
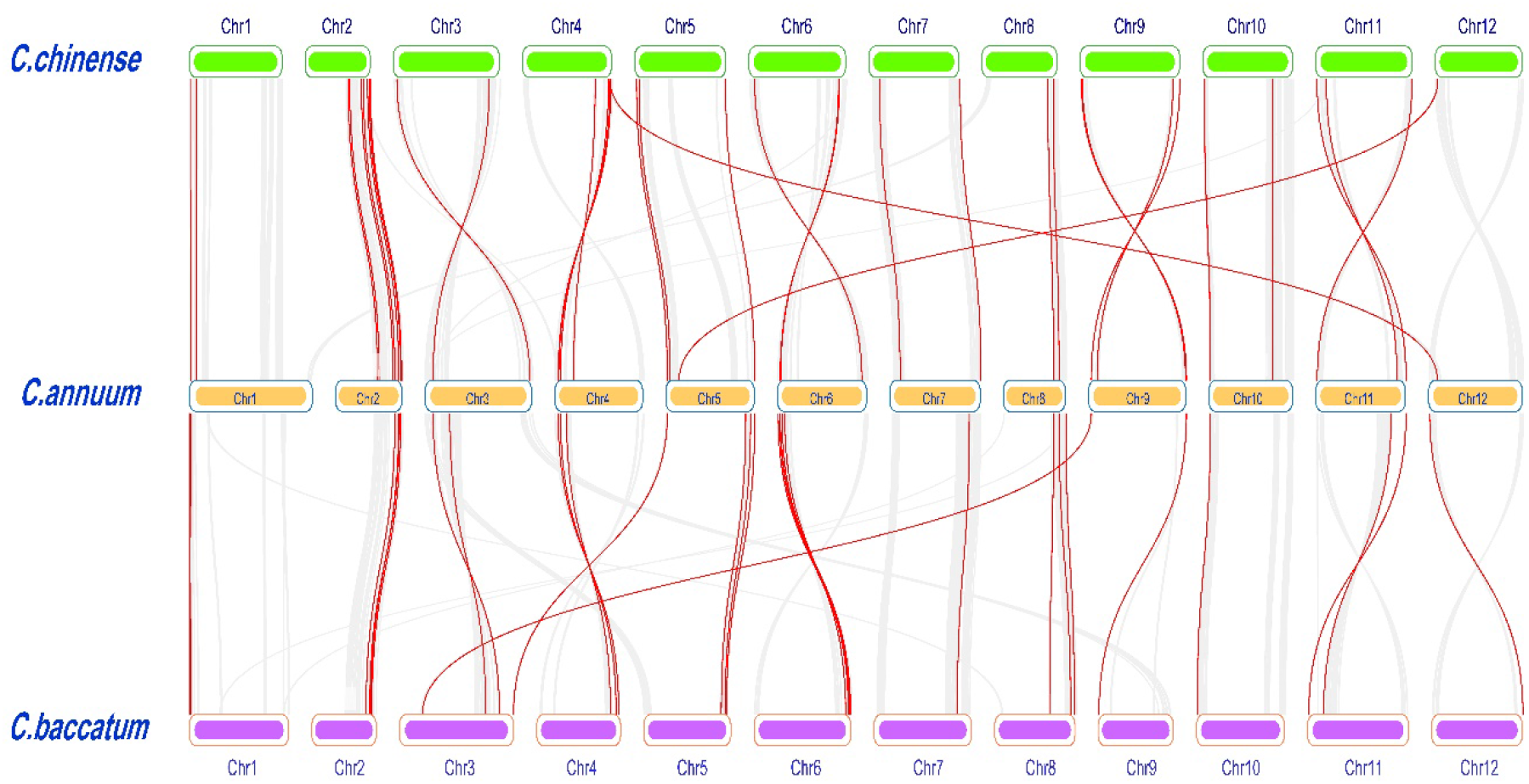
Synteny of the *R2R3-MYB* transcription factors across the *Capsicum* spp. genome. The *Capsicum* R2R3-MYB were mapped to the corresponding chromosomes of *C. chinense*, *C. annuum* and *C. baccatum*. Those with a syntenic relationship are joined by red lines. The gray lines indicate all syntenic blocks in the *Capsicum* spp. genome. Specific gene pairs are listed in Supplemental Table S4. The minimum block size for collinear analysis is set at 30 in MCSanX for interspecies analysis.

### Gene structure analysis of *Capsicum* R2R3-MYBs

Phylogenetic analyses were performed on 108 CaR2R3-MYBs, 106 CbR2R3-MYBs and 110 CcR2R3-MYBs, respectively (Supplementary Figure S1). In addition, a comprehensive phylogenetic analysis was done on 125 *Arabidopsis* R2R3-MYBs, 324 summed R2R3-MYBs in three *Capsicum* spp. and 36 published anthocyanin related MYBs (Table 3, Supplemental Figure S2). The published anthocyanin-related R2R3-MYBs were composed of activators and repressors of anthocyanin biosynthesis (Table 3).

**Table 3.** Plant R2R3-MYB family genes involved in anthocyanin biosynthesis.

| Gene name | Gene ID | Species | Function<br>(activator+/-repressor-) | References |
| --- | --- | --- | --- | --- |
| AmROSEA1 | DQ275529 | *Anitirrhinum majus* | + | Schwinn et al. (2006) |
| AmROSEA2 | DQ275530 | *Anitirrhinum majus* | + | Schwinn et al. (2006) |
| AmVenosa | DQ275531 | *Anitirrhinum majus* | + | Schwinn et al. (2006) |
| AtPAP1/AtMYB75 | AT1G56650 | *Arabidopsis thaliana* | + | Borevitz et al. (2000) |
| AtPAP2/AtMYB90 | AT1G66390 | *Arabidopsis thaliana* | + | Borevitz et al. (2000) |
| AtPAP3/AtMYB113 | AT1G66370 | *Arabidopsis thaliana* | + | Gonzalez et al. (2008) |
| AtPAP4/AtMYB114 | AT1G66380 | *Arabidopsis thaliana* | + | Gonzalez et al. (2008); Heppel et al. (2013) |
| AtMYB60 | AT1G08810 | *Arabidopsis thaliana* | - | Kranz et al. (1998); Park et al. (2008) |
| CaMYBA | AJ608992 | *Capsicum annuum* | + | Borovsky et al. (2004);<br>Aguilar-Barragán and Ochoa-Alejo (2014) |
| FaMYB1 | MN689833 | *Fragaria x ananassa* | - | Aharoni et al. (2001) |
| FaMYB10 | EU155162 | *Fragaria vesca* | + | Lin-Wang et al. (2014) |
| FcMYB1 | GQ867222 | *Fragaria chiloensis* | - | Salvatierra et al. (2013) |
| GhMYB10 | AJ554700 | *Gerbera hybrida* | + | Elomaa et al. (2003) |
| GtMYB3 | AB289445 | *Gentiana triflora* | + | Nakatsuka et al. (2008) |
| IbMYB1 | AB258984 | *Ipomoea batatas* | + | Mano et al. (2007) |
| MdMYB1 | DQ886414 | *Malus domestica* Borkh. | + | Takos et al. (2006) |
| MdMYB3 | JN544704 | *Malus* × *domestica* | + | Vimolmangkang et al. (2013) |
| MdMYB6 | DQ074461 | *Malus* × *domestica* | - | Gao et al. (2011) |
| MdMYB10 | EU518249 | *Malus* × *domestica* | + | Espley et al. (2007) |
| MtLAP1 | FJ199998 | *Medicago truncatula* | + | Peel et al. (2009); Bond et al. (2016) |
| MtMYB2 | AES99346 | *Medicago truncatula* | - | Jun et al. (2015) |
| NtAN2 | FJ472647 | *Nicotiana tabacum* | + | Pattanaik et al. (2010) |
| PacMYBA | JN166079 | *Prunus avium* L | + | Shen et al. (2014) |
| PhAN2 | AAF66727 | *Petunia hybrida* | + | Quattrocchio et al. (1999) |
| PhDPL | HQ116169 | *Petunia hybrida* | + | Albert et al. (2011) |
| PhMYB27 | KF985023 | *Petunia hybrida* | - | Mur (1995); Albert et al. (2011) |
| PhPHZ | HQ116170 | *Petunia hybrida* | + | Albert et al. (2011) |
| PyMYB10 | KF387520 | *Pyrus pyrifolia* | + | Feng et al. (2010) |
| SlANT1<sup>AC</sup>/LeANT1 | AY348870 | *Solanum lycopersicum* cv. *Ailsa Craig* | + | Mathews et al. (2003) |
| SlANT1<sup>Aft</sup> | ABO26065 | *Solanum lycopersicum*.L accession LA1996 | + | Sapir et al. (2008) |
| SlAN2 | FJ705320 | *Solanum lycopersicum*.L | + | Kiferle et al. (2015) |
| SlMYBL2 | XM_004239350 | *The Pro35S:BrTT8 transgenic Solanum lycopersicum* Mill. cv *Ailsa Craig, AC* | - | Zhang et al. (2019) |
| StAN1 | DQ917781 | *Solanum tuberosum* L. | + | Jung et al. (2009) |
| StAN2 | AY841131 | *Solanum tuberosum* L. | + | Jung et al. (2009) |
| StMYBA1 | KP317177 | *Solanum tuberosum* L. | + | Liu et al. (2016); Liu et al. (2017) |
| VvMYB5b | AY899404 | *Vitis vinifera* | + | Deluc et al. (2008); Cavallini et al. (2014) |

The exon-intron structure pattern of the MYBs was further investigated in *C. annuum*, *C. baccatum* and *C. chinense* (Supplementary Figure 2). Generally, over 95% of R2R3-MYBs in *Capsicum spp*. had at least one intron over the entire gene sequence. The majority of R2R3-MYBs in pepper had a structure which was composed of two introns with three exons (respectively 70 of 108 CaMYBs, 69 of 106 CbMYBs and 72 of 110 CcMYBs).

### Identification of R2R3-MYB repressors

In total 53 candidate R2R3-MYB repressors were identified based on their repression motifs in *Capsicum* spp. (Supplementary Figure 2, Table 4). To be specific, 47 candidate R2R3-MYB repressors only had an EAR motif (15 CaMYBs, 15 CbMYBs and 17 CcMYBs), three had a LxLxPP motif (CaMYB37, CbMYB32 and CcMYB40) and three had both EAR- and LxLxPP motifs (CaMYB16, CbMYB13 and CcMYB16).

**Table 4.** Capsicum spp. R2R3-MYB repressors with conserved repressor motifs.

| Repression motif | EAR (LxLxL, DLNxxP) motif | LxLxPP motif |
| --- | --- | --- |
| *Capsicum annuum* | *CaMYB5, CaMYB16, CaMYB34, CaMYB45, CaMYB48, CaMYB50, CaMYB55, CaMYB67, CaMYB68, CaMYB70, CaMYB72, CaMYB82, CaMYB91, CaMYB92, CaMYB93, CaMYB101* | *CaMYB16, CaMYB37* |
| *Capsicum baccatum* | *CbMYB4, CbMYB13, CbMYB29, CbMYB46, CbMYB50, CbMYB66, CbMYB67, CbMYB69, CbMYB76, CbMYB85, CbMYB89, CbMYB92, CbMYB93, CbMYB94, CbMYB102, CbMYB106* | *CbMYB13, CbMYB32* |
| *Capsicum chinense* | *CcMYB5, CcMYB16, CcMYB39, CcMYB47, CcMYB51, CcMYB52, CcMYB61, CcMYB72, CcMYB79, CcMYB87, CcMYB91, CcMYB92, CcMYB93, CcMYB103, CcMYB104, CcMYB105, CcMYB108, CcMYB110* | *CcMYB16, CcMYB40* |

To identify anthocyanin-related R2R3-MYB repressors in pepper, a phylogenetic analysis was performed using candidate pepper MYB repressors together with 36 published anthocyanin MYB regulators from other plant species (Table 3, Figure 4). Meanwhile, CaMYB101, CbMYB89 and CcMYB92 were characterized within one cluster (red curve in Figure 4), together with two identified R2R3-MYB repressors from the *Solanaceae* family (petunia PhMYB27 and tomato SlMYBL2) (Albert et al., 2011; Zhang et al., 2019). In addition, two strawberry R2R3-MYB repressors (FaMYB1 and FcMYB1) and one *Medicago truncatula* R2R3-MYB repressor (MtMYB2) were the closest orthologs to the cluster of CcMYB87, CcMYB108, CbMYB67 and CaMYB70 and the cluster of CaMYB101, CbMYB89 and CcMYB92. Moreover, CaMYB16, CbMYB13 and CcMYB16 were characterized together with an identified apple R2R3-MYB repressor (MdMYB6), while CcMYB91 was characterized together with an identified *Arabidopsis* R2R3-MYB repressor (AtMYB60).

**Figure 4.**
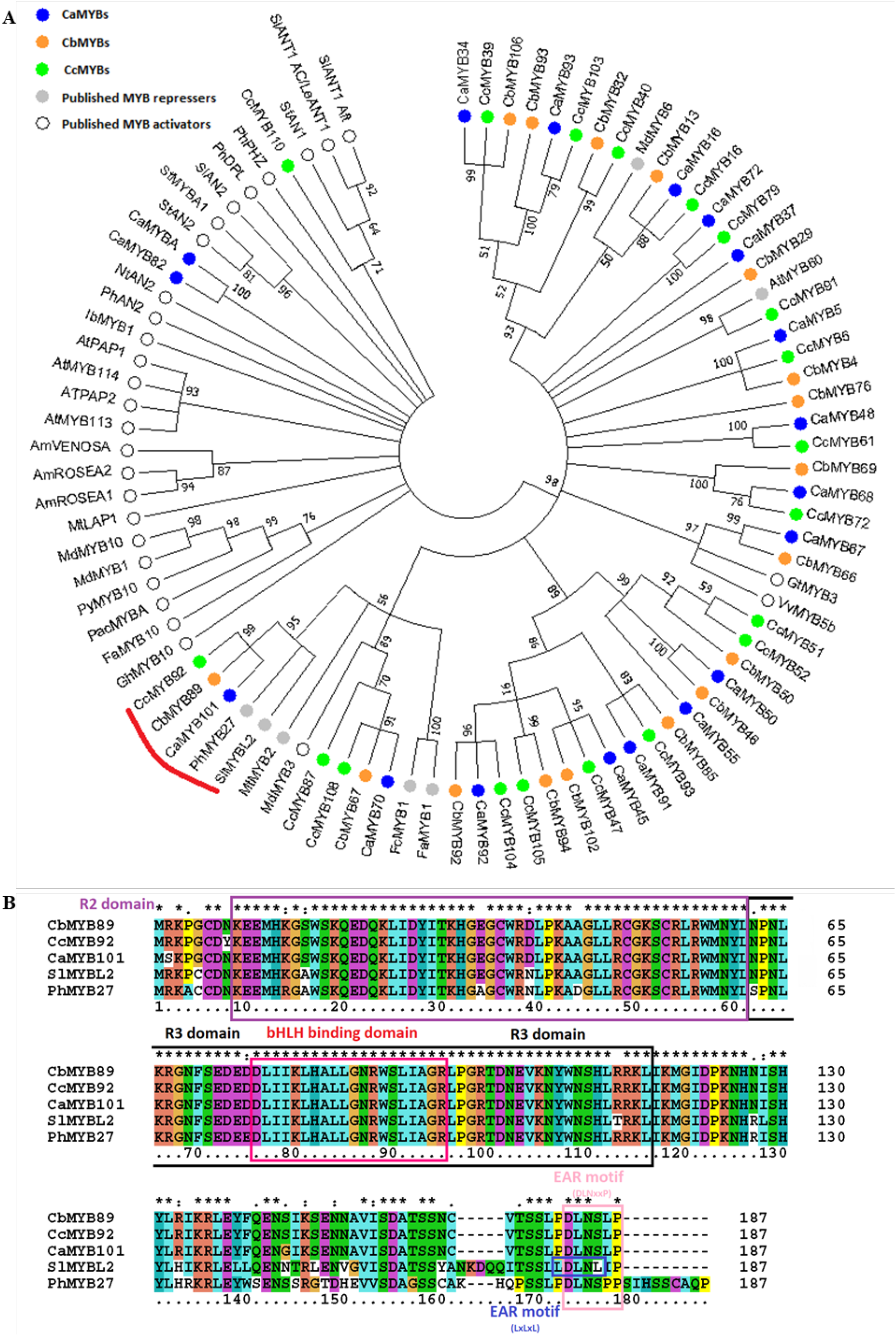
(A)Phylogenetic tree of R2R3-MYB with repression motif and published anthocyanin-related MYBs. CaMYB82 and CaMYB_A_ is the same gene. The red curve shows the cluster of CaMYB101, CbMYB89 and CcMYB89 with petunia and tomato MYB repressors. The blue, orange and green dots indicate CaMYBs, CbMYBs and CcMYBs, respectively (Table 4). The white and grey dots indicate published MYB activators and MYB repressors (Table 3). (B) Multiple alignment of predicted protein sequences of CaMYB101, CbMYB89, CcMYB92, SlMYBL2 and PhMYB27. The R2, R3 domains, EAR motifs (LxLxL and DLNxxP) and bHLH binding domain is labeled above the alignment.

CaMYB101, CbMYB89 and CcMYB92 had high amino acid similarity with other *Solanaceae* family members such as tomato SlMYBL2 and petunia PhMYB27, especially the R2, R3 domain and EAR motif (Figure 4B). It showed that the conserved R2, R3 domain and EAR motif of CaMYB101, CbMYB89 and CcMYB92 were identical. The tomato SlMYBL2 had two EAR motifs, LxLxL and DLNxxP, compared with the other four MYBs with one EAR motif.

### Expression profile of candidate CaR2R3-MYB repressors in different tissues

The expression profile of 17 candidate R2R3-CaMYB repressors (Table 4), based on the RNA-seq transcriptome analysis of *C. annuum* cv. Zunla (Qin et al., 2014), revealed expression variation in different tissues and fruit developmental stages (Figure 5). Eight (CaMYB37, CaMYB45, CaMYB67, CaMYB70 CaMYB72, CaMYB91, CaMYB92 and CaMYB101) out of 17 R2R3-CaMYB repressors showed a relatively high expression level in fruits, especially at pre-breaker stage (3-4cm fruit length) and at post breaker stage (5 days after breaker fruits), respectively (Figure 5).

**Figure 5.**
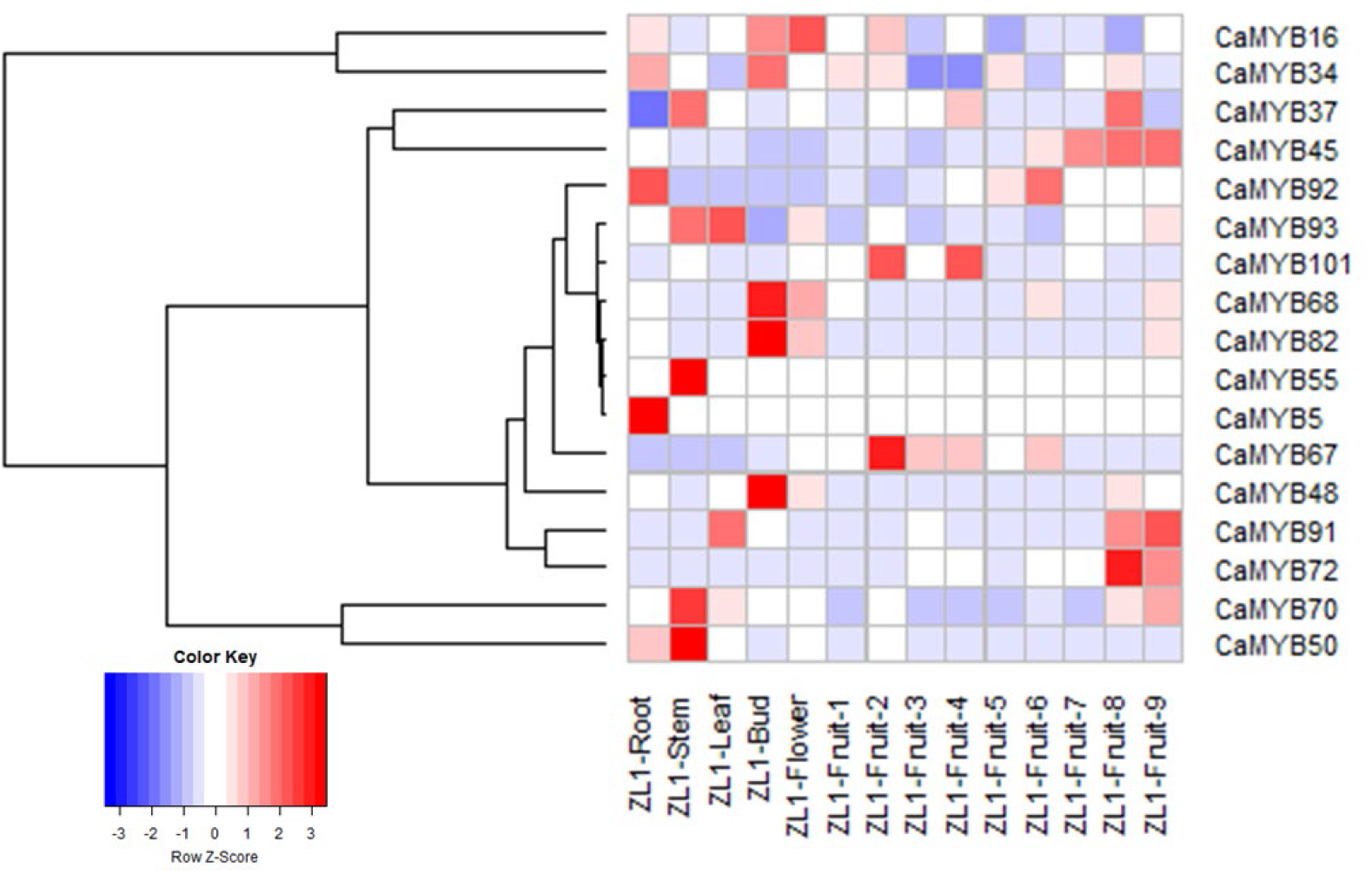
The gene expression patterns in root, stem, leaf, bud, flower and fruits of *Capsicum annuum* cv. Zunla. The color indicates relative gene expression in selected tissues based on rpkm (reads per kb per million mapped reads). Fruit stages 1-3 represent pre-breaker stages (1-3cm, 3-4cm, 4-5cm fruit length; mature green), fruit stages 4-6 represent the breaker stage (when the fruit was turning red) and fruit stages 7-9 represent post-breaker stages (3, 5, and 7 days after breaker).

### Functional analysis of CaMYB101 in *C. annuum* cv. Tequila

The *C. annuum* cv. Tequila is a transiently purple-fruited genotype of which the fruit is green in the early stages of fruit development and then turns purple and eventually red when fully ripe. To explore the involvement of MYB repressors in the regulation of fruit peel color in cv. Tequila, we investigated the candidate gene *CaMYB101*. CaMYB101 is a candidate R2R3-MYB repressor characterized by an EAR motif and, meanwhile, its orthologs in *C. baccatum* (CbMYB89) and *C. chinense* (CcMYB92) were also identified as repressors with an EAR motif (Table 4). Based on the phylogenetic analysis and transcriptome profiling, *CaMYB101* was selected as a candidate gene. CaMYB101 had high homology to tomato SlMYBL2 and petunia PhMYB27 (Figure 4) that are involved in the negative regulation of anthocyanin biosynthesis. The published transcriptome profile also demonstrated a relatively high expression level of *CaMYB101* in pepper fruits (cv. Zunla) (Figure 5), indicating it may function in fruits. In addition, the duplication analysis showed that it did not undergo a duplication event, which minimized the paralog-caused interference and, therefore, functional redundancy. First of all, the expression of *CaMYB101* in purple pepper background, cv. Tequila, was verified by RT-qPCR in different tissues and fruits at different developmental stages (Figure 6). The expression of *CaMYB101* increased from the fertilized ovary to the fruit turning stage (28 DAA) and then decreased until the fruits were fully ripe.

**Figure 6.**
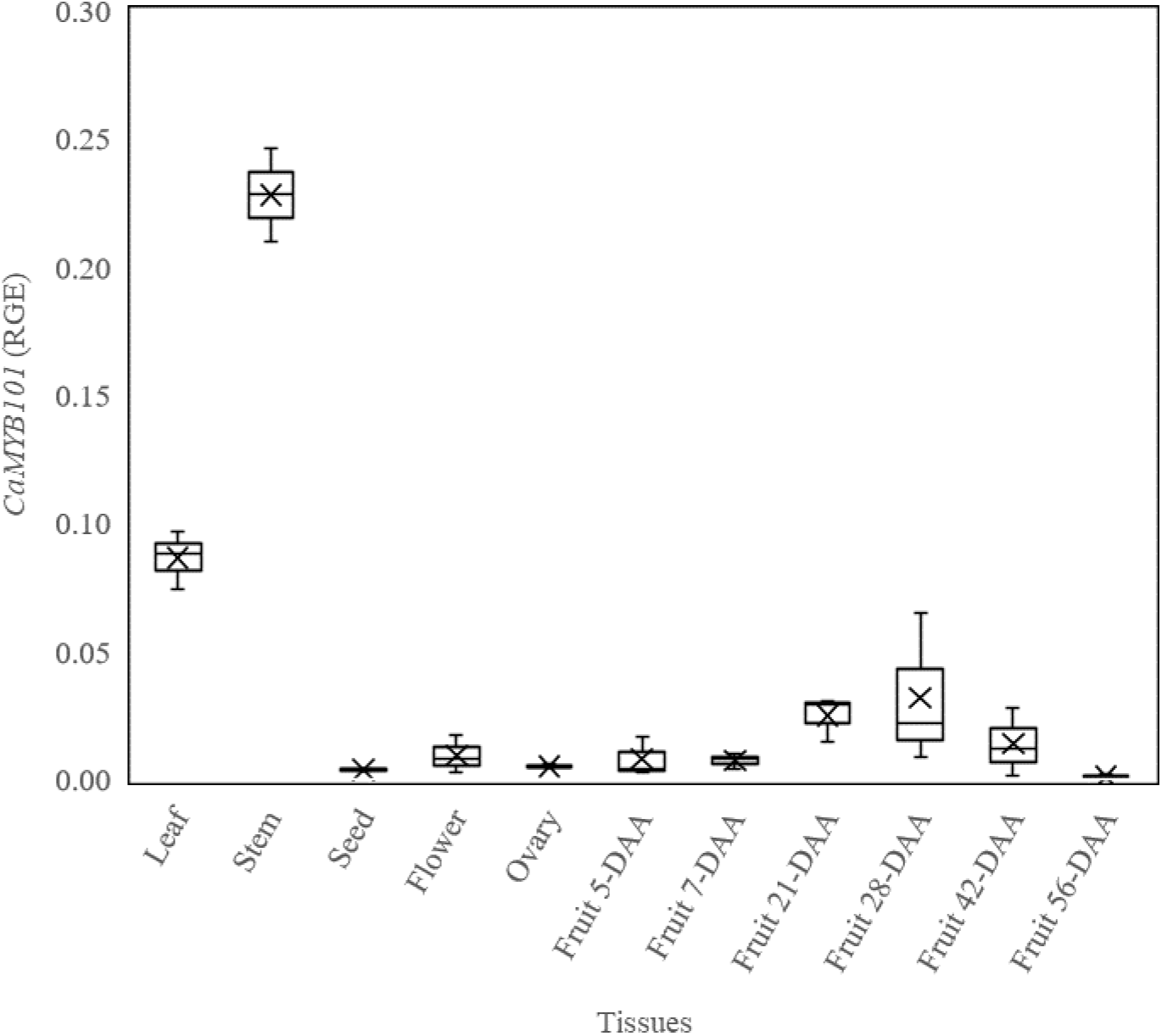
Expression profile of *CaMYB101* genes in different tissues and in fruits during different stages obtained via qRT-PCR. A *Ubiqutin* (*Ub*) gene was used as an internal control. Box plot is generated based on three biological replicates of each tissue (N = 3). Fruit stages refer to Chapter 5 Figure 1.

To investigate the role of CaMYB101 in anthocyanin biosynthesis in pepper fruit, three repeated virus-induced gene silencing (VIGS) experiments were performed with pTRV2::GUS constructs as control. All three VIGS experiments showed the reproducible results. The pTRV2::GUS infected plants produced only green leaves (GUS-GL), while the pTRV2::CaMYB101 infected plants produced both green leaves (MYB-GL) and purple leaves (MYB-PL) that was due to anthocyanin accumulation (Figure 7A). These results indicated that CaMYB101 probably repressed anthocyanin biosynthesis in pepper. To verify the silencing efficiency, the expression of *CaMYB101* was examined in the silenced leaves. Compared with the control plants (GUS-GL), the expression of *CaMYB101* was significantly reduced in leaves of *CaMYB101* silenced plants (in both MYB-GL and MYB-PL) (Figure 7A), which indicated that *CaMYB101* was silenced successfully. Additionally, compared with MYB-GL, the relative expression level of *CaMYB101* was significantly lower in MYB-PL, which further implied the association of *CaMYB101* with anthocyanin accumulation.

**Figure 7.**
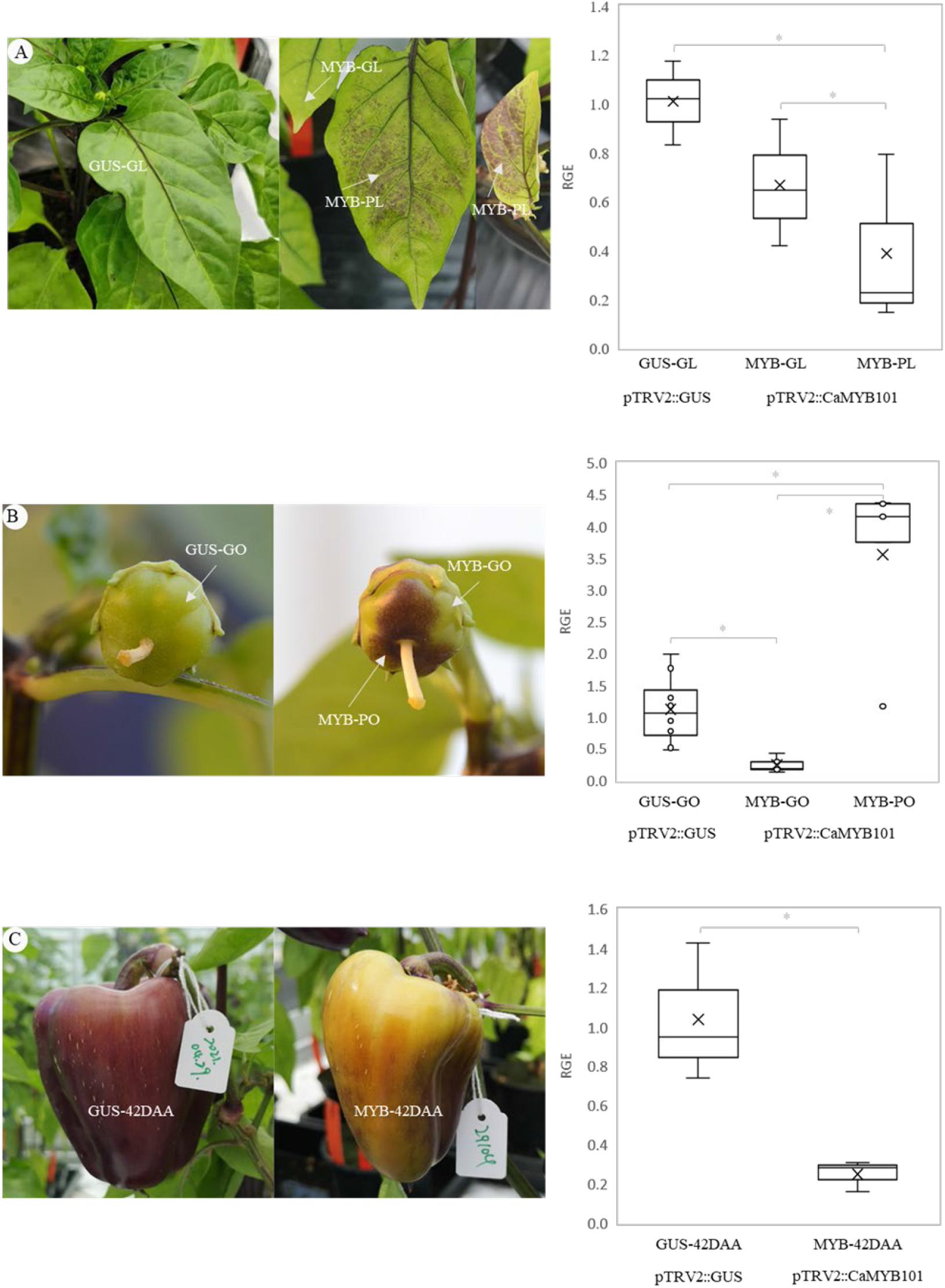
Functional analysis of candidate repressor *CaMYB101* of anthocyanin biosynthesis by VIGS. The VIGS experiments were repeated three times with same phenotypes. Data from the third repetition are presented. Phenotyping of pTRV2::CaMYB101 and pTRV2::GUS plants and relative gene expression level of *CaMYB101* in (A) leaves (N = 3), (B) ovaries (N = 5), and (C) fruits (42 DAA; N = 3). GUS-GL, GUS-GO and GUS-42 DAA represent the pTRV2::GUS green leaves, green ovaries and fruits at 42 DAA, respectively. MYB-GL, MYB-GO represent the green leaves and green parts of ovaries of pTRV2:: CaMYB101 silenced plants. MYB-PL and MYB-PO represent the purple leaves and purple parts of ovaries of pTRV2:: CaMYB101 silenced plants. MYB-42 DAA represents the pTRV2:: CaMYB101 42 DAA fruits.

In the early fruit developmental stages, the ovaries of pTRV2::GUS infected plants were green, while the ovaries of CaMYB101 silenced plants showed intense purple sectors due to anthocyanin accumulation (Figure 7B). However, the CaMYB101 silenced fruits that developed from purple ovaries (42 DAA) showed a faster purple discolouration than the pTRV2::GUS fruits (42 DAA) (Figure 7C). We examined the expression of *CaMYB101* in the silenced ovaries and fruits. The clear boundary of anthocyanin pigmentation in *CaMYB101* silenced ovaries (Figure 7B) enabled precise sampling of the purple parts (MYB-PO) and green parts (MYB-GO). In the green parts of *CaMYB101* silenced ovaries (MYB-GO), the expression of *CaMYB101* was also significantly reduced, compared to the green ovaries of control (GUS-GO) (Figure 7B), suggesting that *CaMYB101* was successfully silenced. However, unexpectedly, *CaMYB101* showed a significantly higher expression in the purple part of *CaMYB101* silenced ovaries (MYB-PO) than in the green part of the same ovaries (MYB-GO) and the green ovaries of control (GUS-GO). At a later fruit developmental stage (42 DAA), consistent with leaves, a significantly lower expression of *CaMYB101* was detected in pTRV2::CaMYB101 fruits than in pTRV2::GUS fruits (Figure 7C), which indicated the successful silencing of CaMYB101 in fruits.

The expression of anthocyanin biosynthetic genes was verified in ovaries of pTRV2::CaMYB101 and pTRV2::GUS infiltrated plants. The majority of tested anthocyanin biosynthetic genes were significantly upregulated upon *CaMYB101* silencing, including regulatory genes *CaMYB_A_* (*CaMYB82*), *CabHLH*, EBGs *CaCHS2*, *CaCHI2*, *CaF3H* and *CaF3’H* as well as LBGs *CaF3’5’H*, *CaDFR*, *CaANS* and *CaUFGT*. These genes all showed significantly higher expression level in MYB-PO compared to MYB-GO and/or GUS-GO (Figure 8).

**Figure 8.**
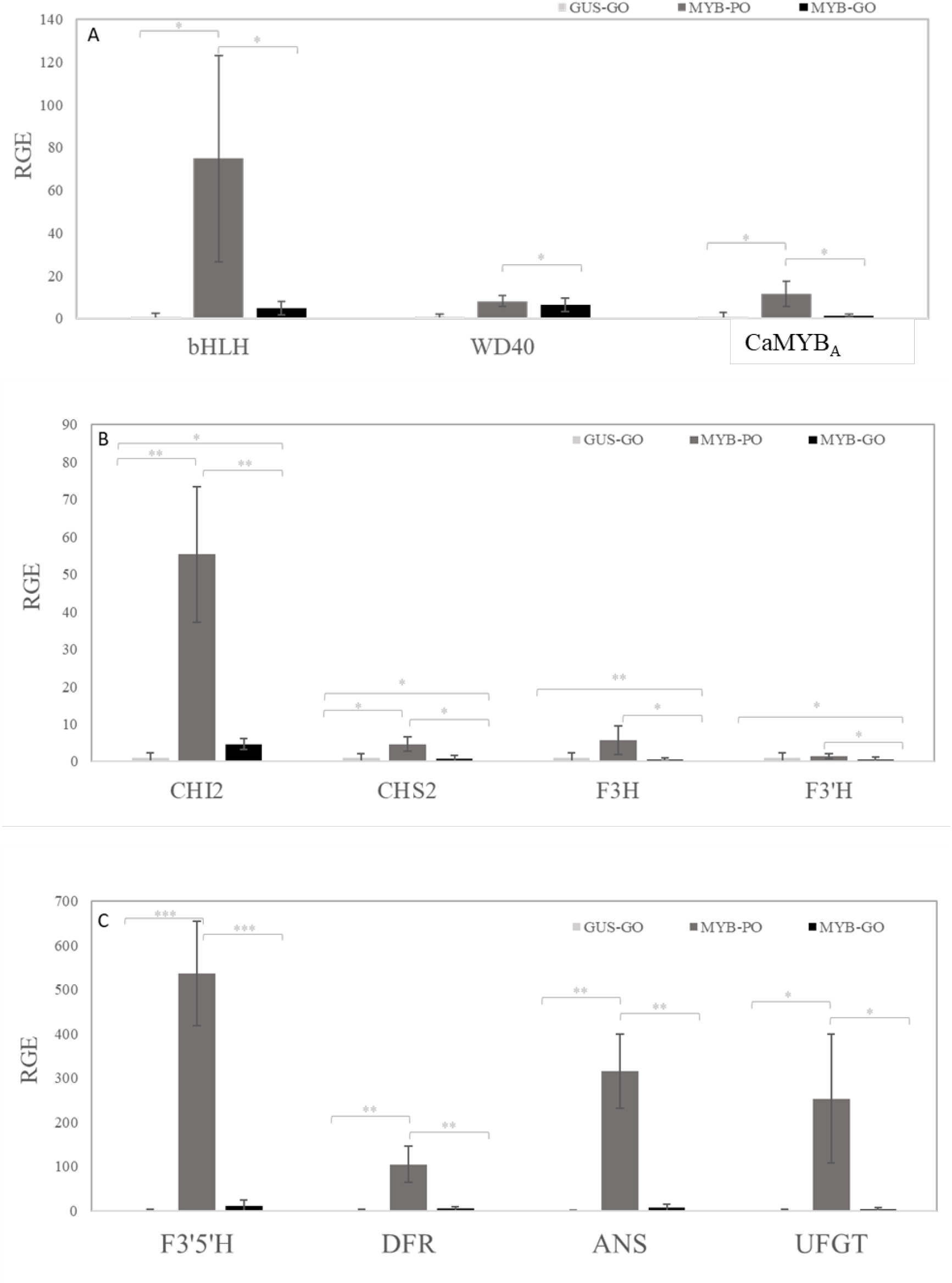
The relative expression level of anthocyanin biosynthetic genes in ovaries (A) *CabHLH, CaWD40* and *CaMYB_A_* activator (CaMYB82 in this study and NCBI accession number AJ608992), (B) anthocyanin early biosynthetic genes *chalcone synthase 2* (CaCHS2), *chalcone isomerase 2 (CaCHI2), flavanone 3-hydroxylase* (CaF3H) and *flavonoid 3’*-*hydroxylase* (CaF3’H) and (C) anthocyanin late biosynthetic genes *flavonoid 3’, 5’*-*hydroxylase* (*CaF3’5’H*), *dihydroflavonol 4*-*reductase* (*CaDFR*), *anthocyanin synthase* (*CaANS*) and a*nthocyanidin 3*-*O*-*glucosyltransferase* (CaUFGT). GUS-GO, MYB-GO and MYB-PO represent the pTRV2::GUS ovaries, purple spots of pTRV2::CaMYB101 ovaries and green parts of pTRV2:: CaMYB101 ovaries, separately.

## Discussion

### Identification and characterization of R2R3-MYBs in three *Capsicum* species

At present, the pepper (*Capsicum*. spp) genome sequences have been released for *C. annuum*, *C. baccatum* and *C. chinense* (Kim et al., 2014; Qin et al., 2014; Kim et al., 2017). In plants, complete and accurate gene annotation enables evolutionary and functional studies of gene families. However, different gene families are not completely annotated in the pepper genus. The MYB family is one of the largest transcription factor families, with the R2R3-MYBs as the main subfamily (Stracke et al., 2001). Nevertheless, there is no comprehensive comparison of R2R3-MYBs across different pepper genomes and most of their functions are unclear. As one of the most cultivated species among the *Capsicum* genus, 108 and 116 *C. annuum* R2R3-MYBs have been genome-widely analyzed based on the Zunla 1.0 genome sequence by Wang et al. (2020) and Arce-Rodríguez et al. (2021), respectively. The reason for the disparity in different number of R2R3-MYBs could be that other known R2R3-MYB plant proteins were also used for identification in Arce-Rodríguez et al. (2021) work. The 108 CaR2R3-MYBs that Wang et al. (2020) identified in *C. annuum* were used for further analysis in this study, together with 106 CbR2R3-MYBs and 110 CcR2R3-MYBs we identified in *C. baccatum* and *C. chinense*, respectively. The number of R2R3-MYBs we identified is similar throughout three *Capsicum* spp., comparable to *Solanum tuberosum* L. (111) (Li et al., 2020), lower than *Solanum lycopersicum* (121) (Zhao et al., 2014) and *Arabidopsis thaliana* (126) (Stracke et al., 2001), and higher than in *Solanum melongena* L. (73) (Wang et al., 2016) and *Oryza sativa* (102) (Yanhui et al., 2006). This suggests that the R2R3-MYBs in different plants evolved at different degrees, indicating that the species with closer evolutionary relationships have a more similar number of R2R3-MYBs.

The genetic conservation, divergence and gene duplication cases of all R2R3-MYBs were studied throughout *Capsicum* spp. including among species (interspecies) and within species (intraspecies). The R2R3-MYB proteins of the three *Capsicum* spp. and *Arabidopsis* shared highly conserved sequences within the R2 and R3 MYB domains based on amino acid frequencies (Figure 1), which confirmed the conserved nature of pepper MYB domains. This is consistent with the previous studies of R2R3-MYBs in Arabidopsis (Dubos et al., 2010). The R2 domain has three conserved tryptophans and the R3 domain has two, where the first tryptophan (missing in R3) was always replaced by a hydrophobic amino acid. These results are consistent with studies in other *Solanaceous* members such as tomato (Zhao et al., 2014), eggplant (Wang et al., 2016) and potato (Li et al., 2020).

Even though the pepper R2R3-MYBs were distributed unevenly on 12 chromosomes within each genome, the overall R2R3-MYB distribution of *C. annuum* and *C. chinense* was more similar compared to *C. baccatum* (Figure 2). In addition, *C. annuum* and *C. chinense* shared more syntenic orthologs (44 pairs) than *C. annuum* and *C. baccatum* (30 pairs) as well as *C. baccatum* and *C. chinense* (21 pairs) (Figure 3, Supplemental Table S4). Both chromosomal distribution and syntenic relationship suggested that *C. annuum* and *C. chinense* were evolutionarily closer, which is in agreement with Kim et al. (2017) who reported that divergence first occurred between *C. baccatum* and the progenitor of *C. annuum* and *C. chinense*. Gene duplication plays an important role in the expansion of the R2R3-MYB family. In addition, the Ka/Ks ratio of over 98 % (117/119) of R2R3-MYBs syntenic gene pairs is smaller than 1.0, which indicates that the R2R3-MYB subfamily underwent purifying selection.

### Characterization of pepper R2R3-MYB repressors

Previous research shows that the R2R3-MYB subfamily plays an important role in anthocyanin synthesis. For example in pepper, an anthocyanin activator, *CaMYB_A_*, has been functionally identified (Borovsky et al., 2004; Zhang et al., 2015). However, no anthocyanin MYB repressor has been identified yet in pepper. We phylogeneticly analysed the predicted pepper R2R3-MYBs with 125 *Arabidopsis* R2R3-MYBs and 34 anthocyanin related R2R3-MYB activators and repressors (Table 3, Supplemental Figure S1) to identify putative anthocyanin-related pepper R2R3-MYBs. The orthologous R2R3-MYBs of *C. annuum, C. baccatum* and *C. chinense* were clustered tightly. Meanwhile, some R2R3-MYBs were clustered with anthocyanin-related R2R3-MYBs. For example, CaMYB101, CbMYB89 and CcMYB92, were within one cluster together with FaMYB1, FcMYB1, MtMYB2, PhMYB27 and SlMYBL2, suggesting that they have similar functions (Figure 4A, Supplemental Figure 1).

The EAR motif is the most predominant transcriptional repression motif in plants, with different sequence patterns, i.e. LxLxL and DLNxxP, and can be used to identify transcriptional repressors (Kagale and Rozwadowski, 2011). In our study, 53 *Capsicum* spp. R2R3-MYBs were identified as repressors by containing at least one EAR motif. A phylogenetic analysis of these 53 candidate repressors and known anthocyanin R2R3-MYB activators and repressors (Table 5A) revealed that (i) CaMYB101, CbMYB89 and CcMYB92 were closely clustered with repressor PhMYB27, SlMYBL2, MtMYB2, FaMYB1 and FcMYB1, (ii) CaMYB16, CbMYB13 and CcMYB16 were clustered together with repressor MdMYB6 and (iii) CcMYB91 was clustered together with repressor AtMYB60 from *Arabidopsis*, indicating their potential function as anthocyanin repressors. Based on this phylogenetic analysis, it was also clear that most candidate repressors had three sets of orthologs in the three pepper genomes and therefore we focused on CaR2R3-MYB repressors, since *C. annuum* is the most widely cultivated species within *Capsicum* spp. The17 CaR2R3-MYB repressors exhibited varied transcript profiles in different tissues according to the public *C. annuum* (Zunla v1.0) RNA-seq data (Figure 5) (Qin et al., 2014). The relative transcript abundance of *CaMYB37*, *CaMYB45*, *CaMYB67*, *CaMYB70*, *CaMYB72*, *CaMYB91*, *CaMYB92* and *CaMYB101* were dependent on the fruit development stage. In purple pepper fruits of cv. Tequila, anthocyanin accumulation is mainly initiated at an early fruit stage (10 DAA) and would degrade during ripening (Liu et al., 2018). The RNAseq data revealed that *CaMYB101*, whose encoded protein sequence clustered with the known anthocyanin MYB repressors PhMYB27 and SlMYBL2, showed high relative-expression levels at the pre-breaker and breaker stages when fruit was turning red. Therefore, *CaMYB101* was selected for further functional study.

### CaMYB101 represses anthocyanin biosynthesis via a negative-feedback loop?

The termination of anthocyanin biosynthesis in purple pepper fruits upon ripening provides us with an opportunity to investigate anthocyanin biosynthesis repression mechanisms. The relative gene expression level of *CaMYB101* in different tissues of cv. Tequila confirmed its expression in fruits (Figure 6). In addition, the expression of *CaMYB101* was strongly associated with purple pigmentation in fruits, which agreed with the expression of *TrMYB133*, an R2R3-MYB repressor for anthocyanin biosynthesis in red forage legumes (Albert, 2015). In addition, the *TrMYB133* as well as *PhMYB27*, the petunia ortholog of *CaMYB101* (Figure 4), both provide feedback repression for anthocyanin biosynthesis through their transcriptional activation by the MBW complexes containing anthocyanin R2R3-MYB activators, and this feedback repression might be conserved across eudicots (Albert et al., 2014; Albert, 2015). Notably, *CaMYB101* had a significantly higher expression level in leaf and stem of cv. Tequila than in fruits, which was different from the RNA-seq profile (Figure 5 & 6). This could be due to the fact that Qin et al., (2014) used a non-purple cultivar, i.e. *C. annuum* cv. Zunla for RNA-seq.

Since anthocyanin R2R3-MYB repressors suppress anthocyanin biosynthesis, we expected that virus-induced silencing of such a repressor would lead to increased anthocynain accumulation by VIGS. Through inoculation of pTRV2::CaMYB101 in cv. Tequila at the cotyledon stage, we found anthocyanin accumulation in both ovaries and leaves, resulting in ovaries and leaves being partially purple and partially green (Figure 7A and 8B). However, VIGS of *CaMYB101* did not result in a more purple fruit phenotype in the later stages of fruit ripening, but instead it accelerated discoloration of purple fruits (Figure 7C). At the same time, *CaMYB101* showed a lower expression in leaves and fruits of pTRV2::CaMYB101 plants compared to pTRV2::GUS plants, confirming the successful silencing of *CaMYB101* in pTRV2::CaMYB101 plants (Figure 7A and 8B). R2R3-MYB repressors suppress anthocyanin biosynthesis via two mechanisms. On the one hand, R2R3-MYB repressors inhibit the assembly of functional MBW complexes by competing with MYB activators to bind to a bHLH transcription factor (LaFountain and Yuan, 2021). On the other hand, they suppress the transcription of anthocyanin structural genes through their repressive motifs. CaMYB101 has a bHLH binding domain and an EAR motif (Figure 4), suggesting it may have both repressive abilities.

In purple areas of pTRV2::CaMYB101 ovaries, the anthocyanin regulator genes, including R2R3-MYB activators *CaMYB_A_*, and most structural genes were activated and showed higher transcript levels than in green parts of the pTRV2::CaMYB101 and pTRV2::GUS ovaries. (Figure 8A-C). Unexpectedly, the expression level of *CaMYB101* in purple areas of pTRV2::CaMYB101 ovaries was also significantly higher compared to green parts of pTRV2::CaMYB101 ovaries, as well as in green pTRV2::GUS ovaries (Figure 7B). The high expression level of *CaMYB101* in the purple area of pTRV2::CaMYB101 ovaries is in line with a study that also reported a high expression level of its tomato orthologous gene *SlMYBL2* in purple fruits of transgenic tomato lines overexpressing *BrTT8* (Zhang et al., 2019), suggesting that *SlMYBL2* was transcriptionally activated to counterbalance the active transcription of the MBW complex to prevent excessive anthocyanin synthesis. Zhang et al. (2019) proposed that an excess of anthocyanin synthesis could act as a signal to activate the expression of *SlMYBL2*. Similarly, *AtMYB4, PhMYB27* and *Tr-MYB133* are direct targets of MBW complexes containing anthocyanin R2R3-MYB activators and they in turn provide negative feedback regulation to MBW complexes (Jin et al., 2000; Albert et al., 2011; Albert, 2015). We hypothesize that anthocyanin accumulation in the purple ovary is due to the initial silencing of *CaMYB101* by VIGS. We further hypothesise that either high anthocyanin levels or high transcription levels of the MBW complex act as a signal to activate *CaMYB101* expression to avoid overproduction of anthocyanins in a negative-feedback loop. It also indicates that the VIGS effect that should be still present in the silenced tissue is not enough to cope with that feedback induction effect.

Even though VIGS of *CaMYB101* caused anthocyanin accumulation in both pepper leaves and ovaries, feedback regulation seemed not to occur in leaves (Figure 7A), suggesting that different molecular mechanisms of anthocyanin pigmentation may exist in vegetative tissues and in ovaries. However, why silencing of *CaMYB101* accelerated anthocyanin discolouration in fruits is still unclear (Figure 7C) and this needs further research. It is recommended to sample more fruit stages during fruit development, especially at an earlier stage to see (i) if the proposed negative feedback loop is also detectable in fruits to overcome the silencing. However, it is worth to realise that fruits probably produce CaMYB101 as part of their “normal” anthocyanin regulation (Figure 6). (ii) if the transient expression of anthocyanin biosynthetic genes in silenced fruits is ahead of time compared to non-silenced fruits. Last but not least, a complete knockout of CaMYB101 would be interesting, since in that case there is no compensation caused by re-activation of this gene.

## Conclusion

The present study provides an overview of the pepper R2R3-MYB gene family by conducting a comprehensive global genome analysis of three *Capsicum*.spp., with specific focus on R2R3-MYB repressors. Functional analysis of *CaMYB101* showed that *CaMYB101* functions as a repressor of anthocyanin accumulation in pepper leaves and ovaries. Our study sheds new light on the negative regulation of anthocyanin biosynthesis in pepper and demonstrates the potential of VIGS to study the function of anthocyanin-related candidate genes in this important horticultural crop. This may lead to novel breeding strategies to modify the anthocyanin content of pepper.

## Methods

### Plant material

Bell pepper (*Capsicum annuum*) cv. Tequila (Enza Zaden) was grown under standard greenhouse conditions in Wageningen, the Netherlands. Nine plants were grown for determining gene expression profile of *CaMYB101* in different plant tissues. Plant leaf, stem, seed, ovary, flower and fruits at day after anthesis (DAA) of 5, 7, 21, 28, 42 and 56 were sampled for tissue specific expression. Tissues and fruits from each three plants were pooled together as one biological replicate.

### Identification of the R2R3-MYB subfamily genes in *Capsicum* spp

Multiple *de novo* pepper genome sequences of *Capsicum baccatum* and *Capsicum chinense* were downloaded from the Pepper Genome Platform (http://peppergenome.snu.ac.kr/). A Hidden Markov Models (HMM) profile of MYB DNA-binding domain (PF00249) was downloaded from Pfam database (http://pfam.xfam.org/). It was used as a query to search the *C. baccatum* and *C. chinense* genomes to identify all MYB containing sequences with an E-values < 1e^−3^. All candidate protein sequences were examined using the NCBI conserved domain database (CDD) (https://www.ncbi.nlm.nih.gov/Structure/cdd/wrpsb.cgi) and ExPASy (https://web.expasy.org/protparam/) to verify the presence of R2 and R3 domains. The R2R3-MYB genes of *C. annuum* used in this study were identified by Wang et al. (2020). The pepper R2R3-MYBs were named based on their species, namely CaMYB for *C. annuum*, CbMYB for *C. baccatum* and CcMYB for *C. chinense*, and numbered based on chromosomal orders (chr1 - chr12, chr0). The amino acid sequences of R2 and R3 MYB repeats of CaMYBs, CbMYBs and CcMYBs were aligned with ClustalW (MEGA-7) and manually adjusted referring to *Arabidopsis* (Dubos et al., 2010). The sequence logos of R2 and R3 MYB repeats were constructed by WebLogo (Crooks et al., 2004). The chromosomal distribution of pepper R2R3-MYBs was mapped using Mapchart 2.2 software (Voorrips, 2002).

### Gene structure motif and phylogenetic analysis

The MEME v5.1.0 online tool was used for the conserved domain investigation. The map of exon-intron structure of *Capsicum* spp. R2R3-MYBs was constructed by TBtools software using coding sequences with the corresponding protein sequences (Chen et al., 2020). The complete protein sequences of *Capsicum* spp. R2R3-MYBs were aligned by ClusterW method then the phylogenetic analysis of R2R3-MYBs was constructed by NJ method with bootstrap test 1000 replicates using MEGA6.0.

### Collinearity/Synteny analyses

The *Capsicum* spp. genome protein sequences and gene structure annotation file were downloaded from Pepper Genome Platform (http://peppergenome.snu.ac.kr/). The Basic Local Alignment Search Tool (BLAST+, NCBI) was used to find the local similarity between sequences with e-value<1e^−5^, number of hits and aligns was set to 5. Then the obtained sequences together with the gene structure annotation file was used to investigate the collinearity/syntenic relationship between *Capsicum* spp. by using a Multiple Collinearity Scan toolkit (Wang et al., 2012). When determining collinearity relationship, the smallest block of intra-species genes was filtered to contain at least 5 pairs and the smallest block of inter-species genes was filtered to contain at least 30 pairs. In this way, the syntenic R2R3-MYB gene pairs were selected. TBtools software was used for the calculation of synonymous (Ks) and non-synonymous (Ka) value.

### Identification of repression motif-containing R2R3-MYB proteins

Repression motifs, included EAR motif (LxLxL, DLNxxP), TLLLFR motif (TLLLFR), R/KLFGV motif (R/KLFGV) and LxLxPP motif (LxLxPP) were screened across all R2R3-MYBs in three pepper genomes (Kagale and Rozwadowski, 2011).

### Expression Profiles of CaMYB Genes Based on RNA-Seq

The expression levels of R2R3-MYBs in *Capsicum annuum* cv. Zunla have been investigated in the different growth development. The RNA-seq atlas was downloaded from China National GeneBank (https://db.cngb.org/search/project/CNPhis0000547/) (Qin et al., 2014). Six different organs including root, stem, leaf, bud, flower and fruits were used for analysis. Furthermore, the relative gene expression level of R2R3=-MYBs were calculated by Reads Per Kilo bases per Million (RPKM). The hierarchical clustering was carried out with Pearson’s correlation distance and the gene clusters were clustered by gene expression.

### VIGS and agrobacteria inoculation

The specific region of CaMYB101 (Capana00g002497) used for silencing was selected by SGN VIGS Tool (Fernandez-Pozo et al., 2015). A 223 bp fragment of CaMYB101 was amplified within the specific region using primers vigsCaMYB101-For (5’- CACCAGGATCTTGGTCTAAACAAGAAGA -3’) and vigsCaMYB101-Rev (5’- CCTATTGCCAAGAAGAGCAT -3’) from cv. Tequila cDNA to design VIGS construct. The fragment was cloned into pTRV2 vector (Liu et al., 2002), then the pTRV2 vectors carrying specific CaMYB101 fragment were transformed into *Agrobacterium tumefaciens* strain GV3101. *A. tumefaciens* culture containing pTRV1, pTRV2::CaMYB101, pTRV2::PDS or pTRV2::GUS was prepared for VIGS experiment as described by Romero et al. (2011). The pTRV1 culture was mixed with pTRV2 culture in a 1:1 ratio. Then TRV infection was done through mixed culture infiltration on the abaxial of cotyledon of 100 three-week-old cv. Tequila seedlings using syringes without a needle. Among them, 45 plants were inoculated with pTRV2::CaMYB101, 45 plants were inoculated with pTRV2::GUS and 10 plants were inoculated with pTRV2::PDS.

### Real-Time Quantitative PCR

The qPCR reaction was prepared with iQ^TM^ SYBR Green Supermix kit (Bio-Rad, USA) then detected by C1000 Touch™ Thermal Cycler (Bio-Rad CFX96 Real-Time system, USA). Relative expression levels of candidate genes were calculated with the 2^−ΔΔCT^ method (Livak and Schmittgen, 2001). Gene specific primers were designed by Primer3Plus (http://www.bioinformatics.nl/cgi-bin/primer3plus/primer3plus.cgi) and the primers used in this paper are listed in Supplemental Table S5.

## Author contribution

YL defined the research question, proposed the methodology and the experimental design. YL and YW carried out the experiments and analysed the experimental data together. ZZ, ZW, SZ and XL performed the bioinformatic analysis. YL and YW wrote the first draft, ZZ revised it. YT, RV, LM, RS and AB provided comments for the final version. All authors have approved the manuscript. This paper has not been accepted or published elsewhere.

