## Supplemental Figures S1-S2 for "Genome-wide characterization of the R2R3-MYB transcription factors in pepper (*Capsicum* spp.) unveils the role of CaMYB101 as repressor in anthocyanin biosynthesis"

**Supplementary Figure 1** Phylogenetic relationships, conserved motifs, and gene structural analysis of R2R3-MYBs in (A) *Capsicum annuum*, (B) *Capsicum baccatum* and (C) *Capsicum chinense*.

**Supplementary Figure 2** Phylogenetic analyses of all *Capsicum* R2R3-MYB transcription factors with Arabidopsis R2R3-MYBs and anthocyanin-related MYB transcription factors from other plant species.

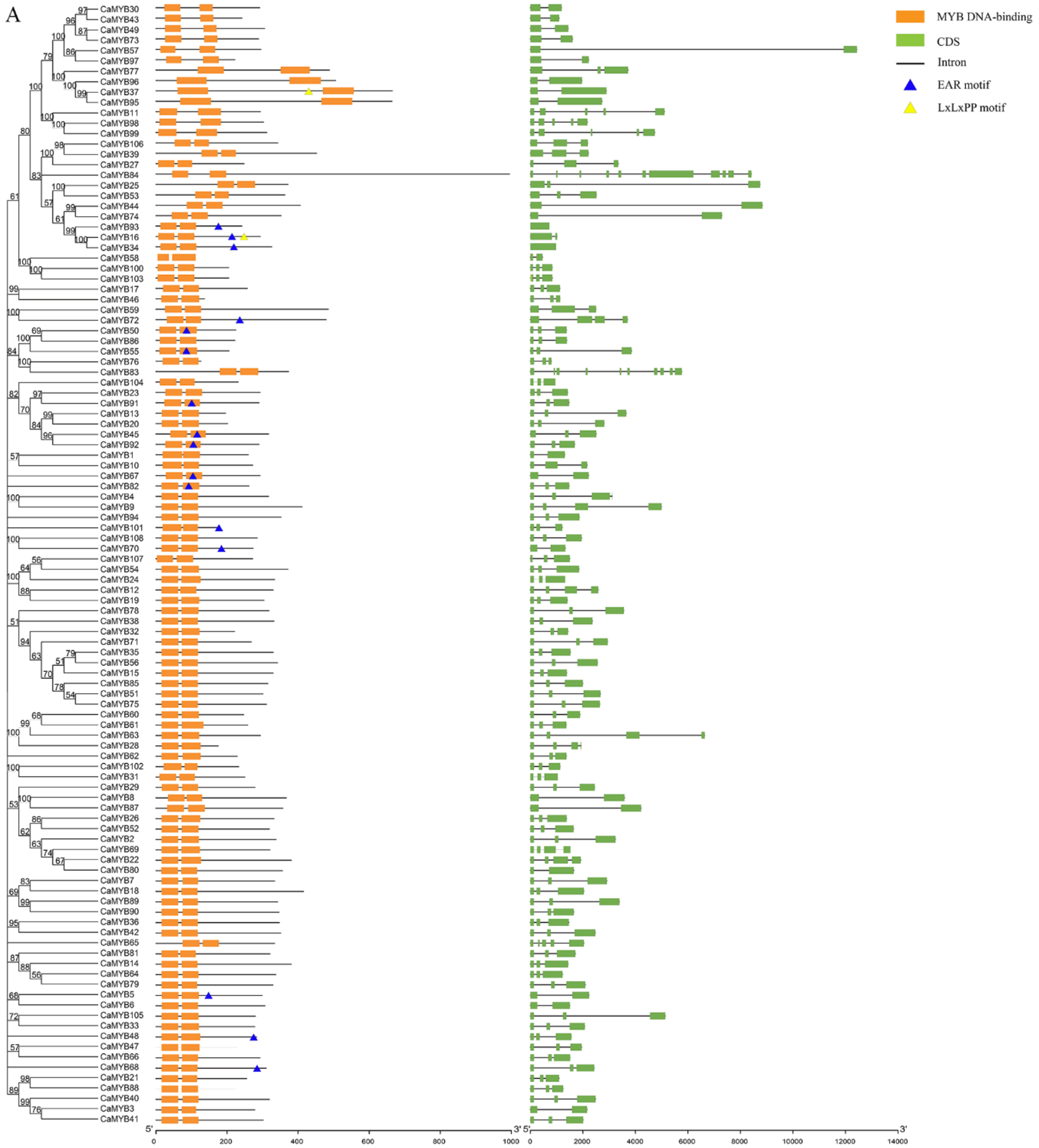

A
MYB DNA-binding
CDS
Intron
EAR motif
LxLxPP motif
CaMYB30
CaMYB43
CaMYB49
CaMYB73
CaMYB57
CaMYB97
CaMYB77
CaMYB96
CaMYB37
CaMYB95
CaMYB11
CaMYB98
CaMYB99
CaMYB106
CaMYB39
CaMYB27
CaMYB84
CaMYB25
CaMYB53
CaMYB44
CaMYB74
CaMYB93
CaMYB16
CaMYB34
CaMYB58
CaMYB100
CaMYB103
CaMYB17
CaMYB46
CaMYB59
CaMYB72
CaMYB50
CaMYB86
CaMYB55
CaMYB76
CaMYB83
CaMYB104
CaMYB23
CaMYB91
CaMYB13
CaMYB20
CaMYB45
CaMYB92
CaMYB1
CaMYB10
CaMYB67
CaMYB82
CaMYB4
CaMYB9
CaMYB94
CaMYB101
CaMYB108
CaMYB70
CaMYB107
CaMYB54
CaMYB24
CaMYB12
CaMYB19
CaMYB78
CaMYB38
CaMYB32
CaMYB71
CaMYB35
CaMYB56
CaMYB15
CaMYB85
CaMYB51
CaMYB75
CaMYB60
CaMYB61
CaMYB63
CaMYB28
CaMYB62
CaMYB102
CaMYB31
CaMYB29
CaMYB8
CaMYB87
CaMYB26
CaMYB52
CaMYB2
CaMYB69
CaMYB22
CaMYB80
CaMYB7
CaMYB18
CaMYB89
CaMYB90
CaMYB36
CaMYB42
CaMYB65
CaMYB81
CaMYB14
CaMYB64
CaMYB79
CaMYB5
CaMYB6
CaMYB105
CaMYB33
CaMYB48
CaMYB47
CaMYB66
CaMYB68
CaMYB21
CaMYB88
CaMYB40
CaMYB3
CaMYB41
5'
3'
0
200
400
600
800
1000
5'
3'
0
2000
4000
6000
8000
10000
12000
14000

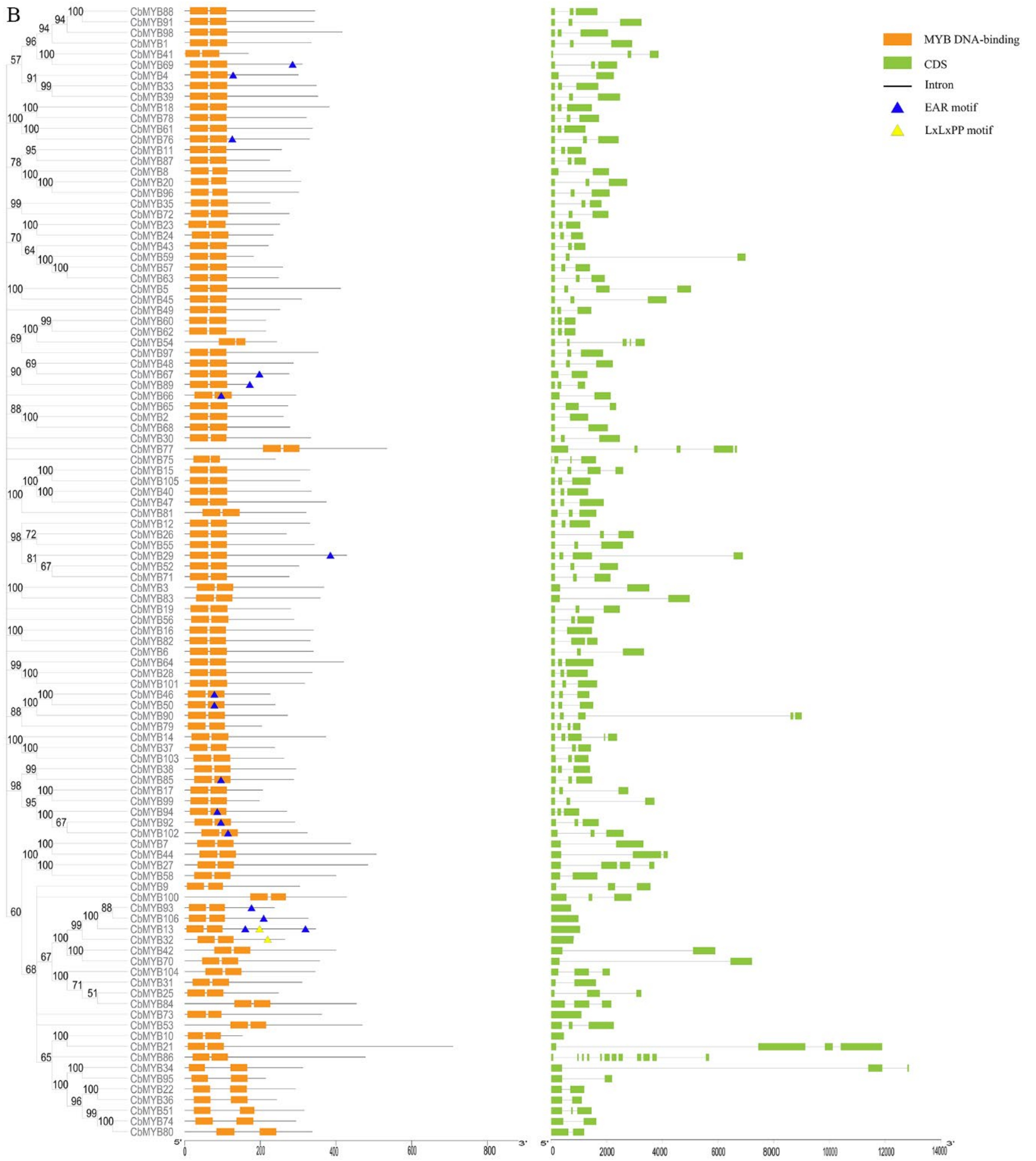
B
MYB DNA-binding
CDS
Intron
EAR motif
LxLxPP motif
CbMYB88
CbMYB91
CbMYB98
CbMYB1
CbMYB41
CbMYB69
CbMYB4
CbMYB33
CbMYB39
CbMYB18
CbMYB78
CbMYB61
CbMYB76
CbMYB11
CbMYB87
CbMYB8
CbMYB20
CbMYB96
CbMYB35
CbMYB72
CbMYB23
CbMYB24
CbMYB43
CbMYB59
CbMYB57
CbMYB63
CbMYB5
CbMYB45
CbMYB49
CbMYB60
CbMYB62
CbMYB54
CbMYB97
CbMYB48
CbMYB67
CbMYB89
CbMYB66
CbMYB65
CbMYB2
CbMYB68
CbMYB30
CbMYB77
CbMYB75
CbMYB15
CbMYB105
CbMYB40
CbMYB47
CbMYB81
CbMYB12
CbMYB26
CbMYB55
CbMYB29
CbMYB52
CbMYB71
CbMYB3
CbMYB83
CbMYB19
CbMYB56
CbMYB16
CbMYB82
CbMYB6
CbMYB64
CbMYB28
CbMYB101
CbMYB46
CbMYB50
CbMYB90
CbMYB79
CbMYB14
CbMYB37
CbMYB103
CbMYB38
CbMYB85
CbMYB17
CbMYB99
CbMYB94
CbMYB92
CbMYB102
CbMYB7
CbMYB44
CbMYB27
CbMYB58
CbMYB9
CbMYB100
CbMYB93
CbMYB106
CbMYB13
CbMYB32
CbMYB42
CbMYB70
CbMYB104
CbMYB31
CbMYB25
CbMYB84
CbMYB73
CbMYB53
CbMYB10
CbMYB21
CbMYB86
CbMYB34
CbMYB95
CbMYB22
CbMYB36
CbMYB51
CbMYB74
CbMYB80
5'
0
200
400
600
800
3'
5'
0
2000
4000
6000
8000
10000
12000
14000
3'

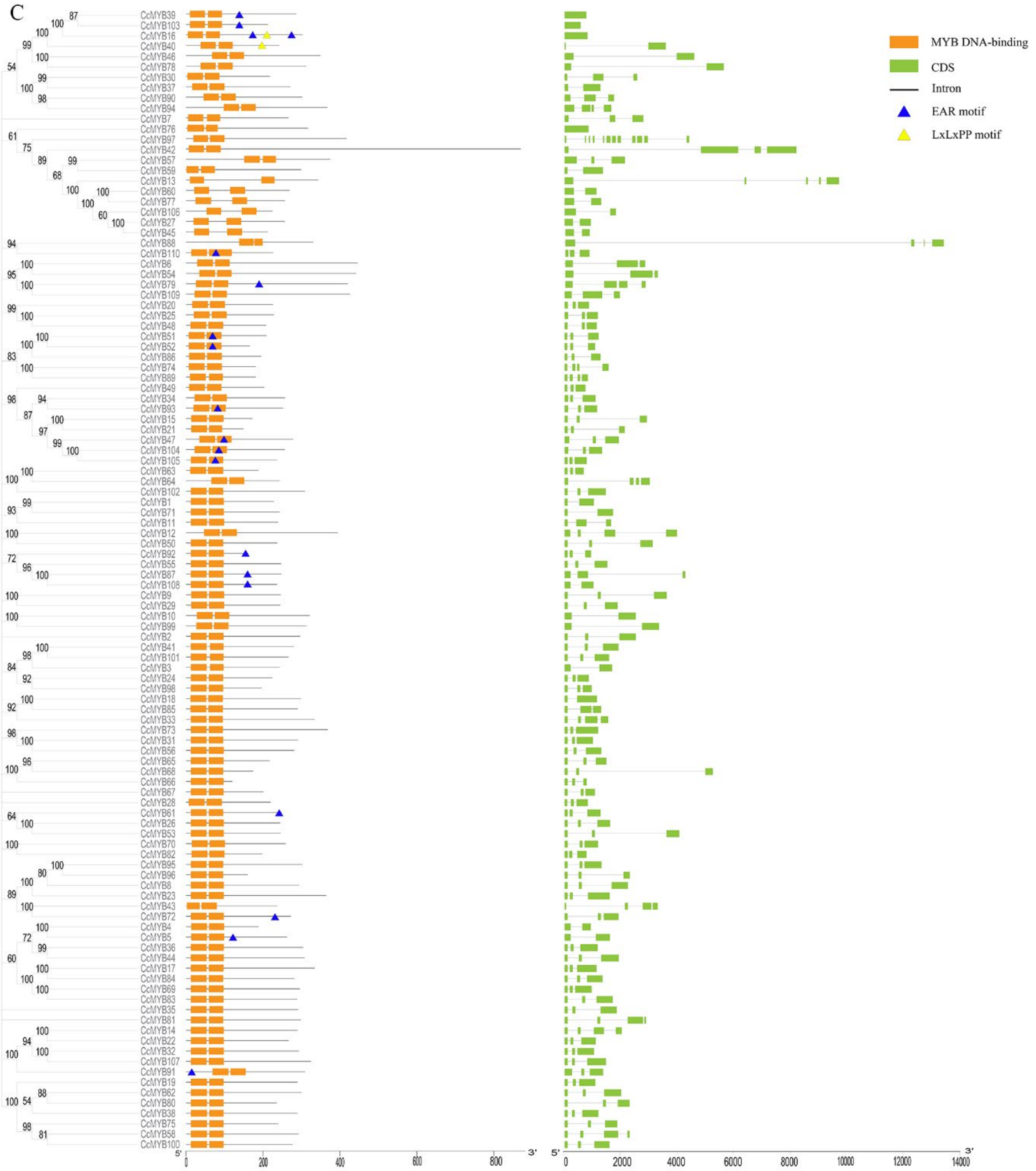


**Supplementary Figure 1.** Phylogenetic relationships, conserved motifs, and gene structural analysis of R2R3-MYBs in (A) *Capsicum annuum*, (B) *Capsicum baccatum* and (C) *Capsicum chinense*. Complete alignments of all *Capsicum* R2R3-MYB proteins were used to construct the phylogenetic tree using the neighbor-joining method. The bootstrap values are indicated on the nodes of the branches. The scale at the bottom provides a size reference. The repression motifs are marked with solid triangles.

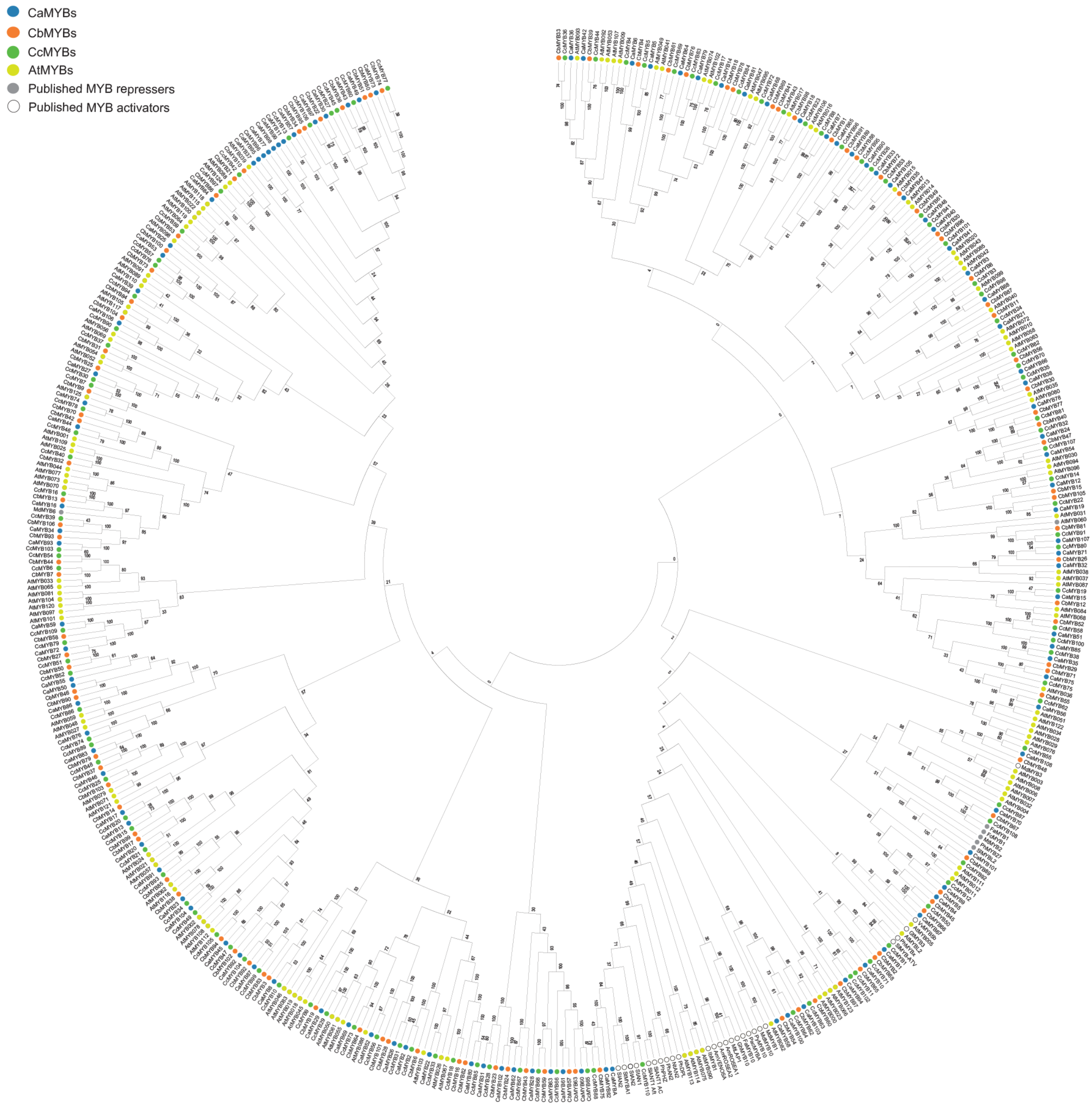


**Supplementary Figure 2.** Phylogenetic analyses of the *Capsicum* R2R3-MYB transcription factors with anthocyanin-related MYB transcription factors from other plant species. The composite phylogenetic tree that included all *Capsicum* and Arabidopsis R2R3-MYB transcription factors and anthocyanin-related MYB transcription factors from Arabidopsis (*Arabidopsis thaliana*), rice (*Oryza sativa*), soybean (*Glycine max*), tomato (*Solanum lycopersicum*), strawberry (*Fragaria x ananassa*), *Gentiana triflora, Ipomoea batatas,* apple *(Malus domestica),* barelclover *(Medicago truncatula),* Tobacco (*Nicotiana tabacum*), Petunia and grape (*Vitis vinifera*) were constructed by the neighbor-Joining method. The bootstrap values are indicated on the nodes of the branches. The anthocyanin-related MYB transcription factors are in bold.
